# Short-Term Plasticity of Contrast Surround Suppression During Repeated Psychophysical Measurement

**DOI:** 10.64898/2026.10.08.757606

**Authors:** Nan Jin, Quan Zhang, Yong-Chun Cai

**Affiliations:** Department of Psychology and Behavioral Sciences, Zhejiang University, Hangzhou, China; Department of Psychology, Ludwig-Maximilians-Universität München, Munich, Germany

**Author notes:** Correspondence concerning this article should be addressed to Yong-Chun Cai, Department of Psychology and Behavioral Sciences, Zhejiang University, 866 Yuhangtang Road, Hangzhou 310058, China.

**Keywords:** surround suppression, perceptual plasticity, divisive normalization, psychophysics

## Abstract

Perceptual surround suppression is commonly treated as stable within a testing session, but recent evidence for normalization reweighting suggests that suppressive interactions can be modified by recent visual input. Here, we asked whether surround suppression changes over minutes of repeated measurement and whether such change varies with stimulus and sampling conditions. Across four psychophysical experiments, we tracked contrast and motion surround suppression while varying center-surround orientation and stimulus spatiotemporal properties. With static gratings, contrast surround suppression increased early and then approached a plateau, with no evidence that this change depended on center-surround orientation. With dynamic gratings, an increase emerged only under finer temporal sampling and more sustained stimulation, and model-based analyses suggested a faster increase than with static stimulation. Motion surround suppression measured with drifting gratings showed no systematic change. Together, these findings show that contrast surround suppression can change within a single psychophysical session without a separate adaptation phase, and that its measured time course varies with stimulus and sampling conditions. The results are consistent with rapid, experience-dependent reweighting of contextual normalization.

## Introduction

Visual encoding of a local stimulus depends not only on the properties of the stimulus itself but also on the surrounding spatial context (Angelucci et al., 2017). Perceptual surround suppression is a prominent example of such contextual modulation and generally refers to conditions in which the presence of surrounding stimuli impairs perception of a central stimulus (Li et al., 2023). One well-studied form is contrast surround suppression: Relative to a central grating presented alone, adding a high-contrast surround reduces the apparent contrast of the central grating or decreases sensitivity to its contrast (Xing & Heeger, 2001; Zenger-Landolt & Heeger, 2003). A related phenomenon in motion perception is motion surround suppression, also termed spatial suppression (Tadin, 2015). At high contrast, observers typically require longer presentation durations to discriminate the motion direction of large stimuli than of small stimuli (Schallmo et al., 2018; Tadin et al., 2003). Although these effects are measured with different stimuli and behavioral tasks, both illustrate how the visual system integrates information over space to regulate local visual representations.

Perceptual surround suppression is generally thought to be closely related to neural surround suppression in the visual cortex, whereby stimuli outside a neuron’s classical receptive field (CRF) modulate, and typically suppress, responses to stimuli within the CRF (Allman et al., 1985; Jones et al., 2001). In humans, the psychophysical magnitude of contrast surround suppression shows a close quantitative correspondence with response suppression in V1, making V1 an important candidate substrate for the perceptual effect (Zenger-Landolt & Heeger, 2003). Motion surround suppression, in turn, has been closely linked to response changes in motion-processing regions, including hMT+ (Schallmo et al., 2018; Tadin et al., 2011).

Despite differences in stimulus configuration, behavioral measure, and associated visual areas, both contrast surround suppression and motion surround suppression can be described within the computational framework of divisive normalization (Cavanaugh et al., 2002; Schallmo et al., 2018; Xing & Heeger, 2001). In this framework, the output of a target neuron or neural population is driven by stimulus-evoked excitation and divisively regulated by activity in a broader normalization pool (Carandini & Heeger, 2012). Applied to visual surround suppression, the central stimulus primarily provides excitatory drive, whereas the surround increases activity in the normalization pool, thereby reducing the response to the center (Carandini & Heeger, 2012; Schallmo et al., 2018). The relative influence exerted by different neurons or neural populations within the normalization pool is commonly expressed by normalization weights. These weights are functional parameters of the computational model and need not correspond one-to-one with specific inhibitory synapses (Carandini & Heeger, 2012; Sato et al., 2016).

Classical normalization models, and many of their standard fitting implementations, treat normalization weights as relatively fixed within the conditions being modeled (Carandini & Heeger, 2012). The Hebbian normalization model proposed by Westrick et al. (2016), however, suggests that these weights can be adjusted dynamically according to the recent joint statistics of sensory inputs. In the model, normalization weights between pairs of neural populations are updated according to a Hebbian rule based on their recent coactivation, with the goal of maintaining a stable product of their paired responses. When two populations frequently respond together, the normalization between them therefore strengthens. This stronger divisive interaction subsequently reduces the extent to which they respond together. Thus, the weight update itself is Hebbian, whereas its functional consequence is anti-Hebbian in the sense that it reduces correlated responses. Therefore, a central prediction from the model is that if contextual suppression depends on adjustable rather than fixed normalization weights, its strength should vary with the recent co-occurrence statistics of the relevant stimulus representations rather than remain a fixed property (Aschner et al., 2018; Yiltiz et al., 2020).

Evidence from animal neurophysiology, human psychophysics, and human neuroimaging provides initial convergence for this prediction (Aschner et al., 2018; Baker et al., 2023; Yiltiz et al., 2020). Cross-orientation suppression (COS), a classic normalization phenomenon in the visual cortex, provides a useful paradigm for testing normalization reweighting. COS refers to the reduction in neural responses or perceptual sensitivity to a target grating when it is superimposed with a differently oriented mask (Bonds, 1989; Morrone et al., 1982). Like surround suppression, COS is also well described by normalization models (Carandini et al., 1997; Carandini & Heeger, 2012). Recording from macaque V1, Aschner et al. (2018) found that adapting to two orthogonal gratings presented synchronously increased subsequent COS, whereas presenting the same gratings asynchronously reduced COS. This suggests that normalization-related suppression can be adjusted according to the recent temporal contingency between inputs. Yiltiz et al. (2020) subsequently observed a similar pattern in human psychophysics. They found that synchronous adaptation to two overlapping gratings produced stronger subsequent COS than alternating adaptation, and the Hebbian normalization model captured this pattern of change. Baker et al. (2023) further examined the possible timescale of normalization reweighting. Using orthogonal target and mask gratings flickering at different frequencies, they tracked suppression over time with steady-state visual evoked responses. EEG and source-localized MEG showed that the increase in COS emerged primarily within approximately 2–5 s after the onset of sustained stimulation. Together, these findings suggest that the strength of visual suppression is not entirely fixed but can be adjusted by the recent joint and temporal statistics of related inputs, as predicted by the Hebbian normalization model, supporting the view of normalization reweighting.

If visual surround suppression depends on normalization and normalization weights are themselves plastic, surround suppression should likewise vary with short-term stimulus history. Theoretically, repeated synchronous presentation of center and surround should strengthen the normalizing influence of surround-related activity on responses to the center and thereby increase perceptual surround suppression. Yiltiz et al. (2020) provided initial support for this prediction. Using an adaptation paradigm, they found stronger contrast surround suppression when the center-surround pairing in the test phase matched the pairing experienced during the adaptation phase, and the change was well described by the Hebbian normalization model. Nevertheless, several questions about short-term changes in perceptual surround suppression remain unresolved. First, previous studies have typically compared suppression before versus after adaptation, or across different adaptation histories, rather than tracking its evolution during continuous behavioral measurement. It is therefore unclear whether experience-dependent changes can be observed within a single session on the timescale of minutes, or how such changes develop and stabilize over time. Second, the feature relationship between center and surround strongly affects the overall magnitude of contrast surround suppression (Schallmo & Murray, 2016; Xing & Heeger, 2001), but it is not known whether this feature selectivity also constrains short-term changes in suppression. Finally, if dynamic changes in contrast surround suppression reflect a more general process of contextual normalization reweighting, an important boundary question is whether a comparable effect extends to motion surround suppression measured with different stimuli and behavioral indices (Tadin et al., 2003).

Across four experiments, we examined whether perceptual surround suppression changes over the course of continuous measurement and probed the boundary conditions of such changes. Experiment 1 used static center-surround gratings and found that contrast surround suppression increased with repeated measurement, with no evidence that the increase depended on the relative orientation of center and surround. Experiment 2 used drifting gratings and found no systematic change in motion surround suppression with repeated measurement. Experiments 3 and 4 then examined whether the spatiotemporal properties of the stimulus modulate changes in contrast surround suppression. Comparing static, high-spatial-frequency stimuli with dynamic, low-spatial-frequency counterphase-flickering stimuli, these experiments showed that contrast surround suppression also increased under dynamic stimulation, but appeared to do so more rapidly. Overall, the findings indicate that perceptual surround suppression can change with recent stimulus history within a single testing session, while the observability and time course of this short-term plasticity depend on stimulus and measurement conditions.

## Experiment 1

Experiment 1 tested whether contrast surround suppression changes with short-term center-surround stimulus input history and whether any such change is modulated by the relative orientation relationship of the center and surround. We predicted that contrast surround suppression would strengthen during repeated measurement containing repeated center-surround stimulation, whereas we made no directional prediction regarding a possible modulation of this increase by center-surround orientation relationship.

### Method

#### Participants

Given that previous studies related to surround suppression, using adaptation paradigms, have reported relatively large changes in suppression (Schallmo & Murray, 2016; Yiltiz et al., 2020), sample-size planning was based on repeated-measures ANOVA with seven time points (see Data Analysis), assuming a large effect size (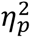 = .14), *α* = .05, and power (1 − *β*) = .80. This yielded a minimum sample size of 8 participants. We recruited 12 students from Zhejiang University, including one author. All participants except the author were naive to the purpose of the experiment. All 12 completed both experimental conditions and were included in the final analysis (6 women, 6 men; age range = 20–24 years, *M* = 21.7, *SD* = 1.7). Participants provided written informed consent and received either monetary compensation or course credit. All procedures were approved by the Research Ethics Committee of Zhejiang University.

#### Apparatus and Stimuli

Visual stimuli were generated in MATLAB R2014a using Psychtoolbox (Brainard, 1997; Pelli, 1997) and presented on a 32-in. Display++ monitor (Cambridge Research Systems Ltd., Rochester, UK; resolution = 1,920 × 1,080 pixels; refresh rate = 100 Hz). The monitor was automatically gamma-linearized. Mean background luminance was 40.26 cd/m². Participants viewed the stimuli binocularly in a dimly lit room at a fixed viewing distance of 75 cm, maintained with a chinrest.

Experiment 1 used two types of static stimuli (Figure 1A). The standard stimulus was a center-surround grating comprising a circular sinusoidal grating at the center (diameter = 1.5° of visual angle; spatial frequency = 4 cpd) surrounded by an annular sinusoidal grating (outer diameter = 6°; spatial frequency = 4 cpd), with a 0.25° gap between center and surround. The outermost 0.25° of the central grating was tapered with a cosine envelope. The Michelson contrasts of the center and surround gratings were fixed at 0.30 and 0.80, respectively, and the two gratings had the same spatial phase. The matching stimulus consisted of the central circular grating presented alone, with the same size, spatial frequency, and edge taper as the central component of the standard stimulus. Its contrast was controlled by an adaptive staircase. On each trial, the orientations of the gratings were randomly set to 45° clockwise or counterclockwise from vertical. Two center-surround orientation conditions were tested. In the parallel condition, center and surround had the same orientation, while in the orthogonal condition, their orientations differed by 90°. Both stimulus types were presented with 100-ms cosine ramps at onset and offset.

**Figure 1.**
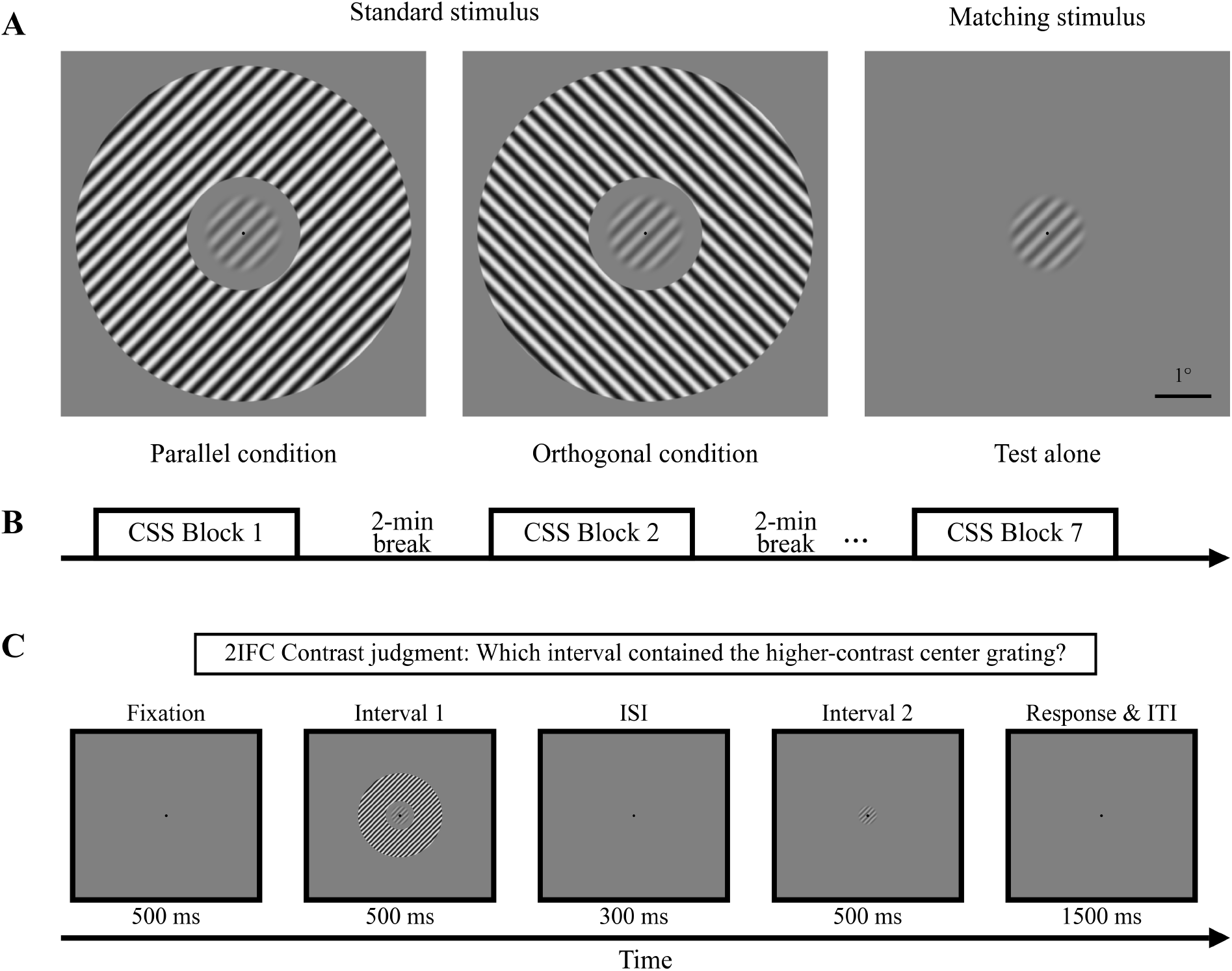
Stimuli and Procedure in Experiment 1. *Note.* (A) Standard and matching stimuli. The standard stimulus consisted of a central grating surrounded by an annular grating. The center and surround had the same orientation in the parallel condition and differed by 90° in the orthogonal condition. The matching stimulus consisted of the central grating presented alone, with its contrast controlled by an adaptive staircase. (B) Session structure in Experiment 1. The parallel and orthogonal conditions were completed on separate days. Each session contained seven 160-trial measurement blocks. (C) Trial sequence. The standard and matching stimuli were presented in random order across two 500-ms intervals. Participants reported which interval contained the central grating with the higher apparent contrast. 2IFC = two-interval forced choice. Stimuli are shown for illustration and are not drawn to scale.

#### Procedure

The parallel and orthogonal conditions were completed on separate testing days at least 24 hr apart, with condition order counterbalanced across participants. As shown in Figure 1B, each testing day comprised seven measurement blocks of 160 trials. Participants took a 30-s break after the 80th trial within each block and a 2-min break between blocks. During all breaks, they were instructed to close their eyes to minimize additional visual input. Each condition comprised 1,120 trials and lasted approximately 1.5 hr. On the day before formal testing, participants completed practice for both stimulus conditions. Practice data were not analyzed.

Contrast surround suppression was measured with a widely used contrast-matching paradigm implemented as a two-interval forced-choice (2IFC) task (Figure 1C) (Xing & Heeger, 2000). Each trial began with a 500-ms fixation display, followed by two 500-ms stimulus intervals separated by a 300-ms fixation-only interval. The standard and matching stimuli each appeared once, in random order. Participants were instructed to ignore the surround, compare the apparent contrast of the central grating across the two intervals, and indicate by keypress whether the center appeared higher in contrast in the first or second interval. No accuracy feedback was provided. To keep trial timing fixed, the response period and intertrial interval together lasted 1.5 s. Within each block, the contrast of the matching stimulus was controlled by two interleaved 1-up-1-down staircases of 80 trials each that started at contrasts of 0.15 and 0.35 and were randomly interleaved. The contrast level of the matching stimulus was selected from 12 logarithmically spaced levels between 0.15 and 0.35. When the matching stimulus was judged higher in contrast, its contrast was reduced by one level the next time that staircase was sampled. When the center of the standard stimulus was judged higher in contrast, matching contrast was increased by one level. Contrast surround suppression was then estimated by fitting a psychometric function to all trials within each block.

#### Data Analysis

For each repeated measurement block in each orientation condition for every participant, the 160 trials from the two staircases were pooled for psychometric fitting. At each matching-stimulus contrast level, we calculated the proportion of trials on which the matching stimulus was judged to have the higher contrast. These response proportions were then fitted in MATLAB using fitnlm with the following cumulative Gaussian function:

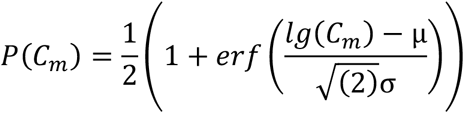

Here, *μ* and *σ* denote the location and scale parameters, respectively, of the cumulative Gaussian function on the log-contrast axis. The matching-stimulus contrast at which the fitted response probability equaled .50 was defined as the point of subjective equality (PSE).

We then defined the Suppression Index (SI) as lg(C₀/PSE), where C₀ is the contrast of the central component of the standard stimulus (0.30). Thus, PSE < C₀ yields SI > 0, indicating that the surrounded central grating was matched to an isolated grating of lower physical contrast and therefore that contrast surround suppression was present. Thus, larger SI values indicate stronger contrast surround suppression.

To test whether suppression increased across repeated measurement blocks, and whether any change depended on the center-surround orientation relationship, SI was submitted to a two-way repeated-measures ANOVA with Orientation Relationship (parallel, orthogonal) and Block (1–7) as within-participant factors. Sphericity was assessed with Mauchly’s test, and Greenhouse-Geisser corrections were applied when the assumption was violated. Bonferroni correction was used for all post hoc comparisons. All statistical analyses were conducted in JASP 0.95.4 (JASP Team, 2025).

#### Results

Figure 2 shows PSE and SI across repeated measurement blocks in the parallel and orthogonal conditions. SI was submitted to a two-way repeated-measures ANOVA with Orientation Relationship (parallel, orthogonal) and Block (1–7) as within-participant factors. The results first revealed a significant main effect of Orientation Relationship, *F*(1, 11) = 17.75, *p* = .001, 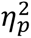 = .62. Averaged across blocks, SI was significantly higher in the parallel condition (*M* = .15, *SD* = .037) than in the orthogonal condition (*M* = .11, *SD* = .036), *t*(11) = 4.21, *p* = .001, Cohen’s *d*=.73, indicating stronger overall contrast surround suppression when the center and the surround shared the same orientation. The main effect of Block was also significant, *F*(1.61, 17.73) = 8.66, *p* = .004, 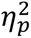 = .44. Bonferroni-corrected post hoc comparisons showed that SI in Block 1 was lower than in Block 2 (adjusted *p* = .006) and in Blocks 3–6 (all adjusted *p*s < .05). This suggested that SI increased after the first block and remained elevated across most subsequent blocks rather than increasing linearly throughout all seven blocks. The Orientation Relationship × Block interaction was not significant, *F*(1.36, 15.00) = .65, *p* = .48, 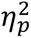 = .06. We therefore found no evidence that the temporal profile of SI differed between the parallel and orthogonal conditions.

**Figure 2.**
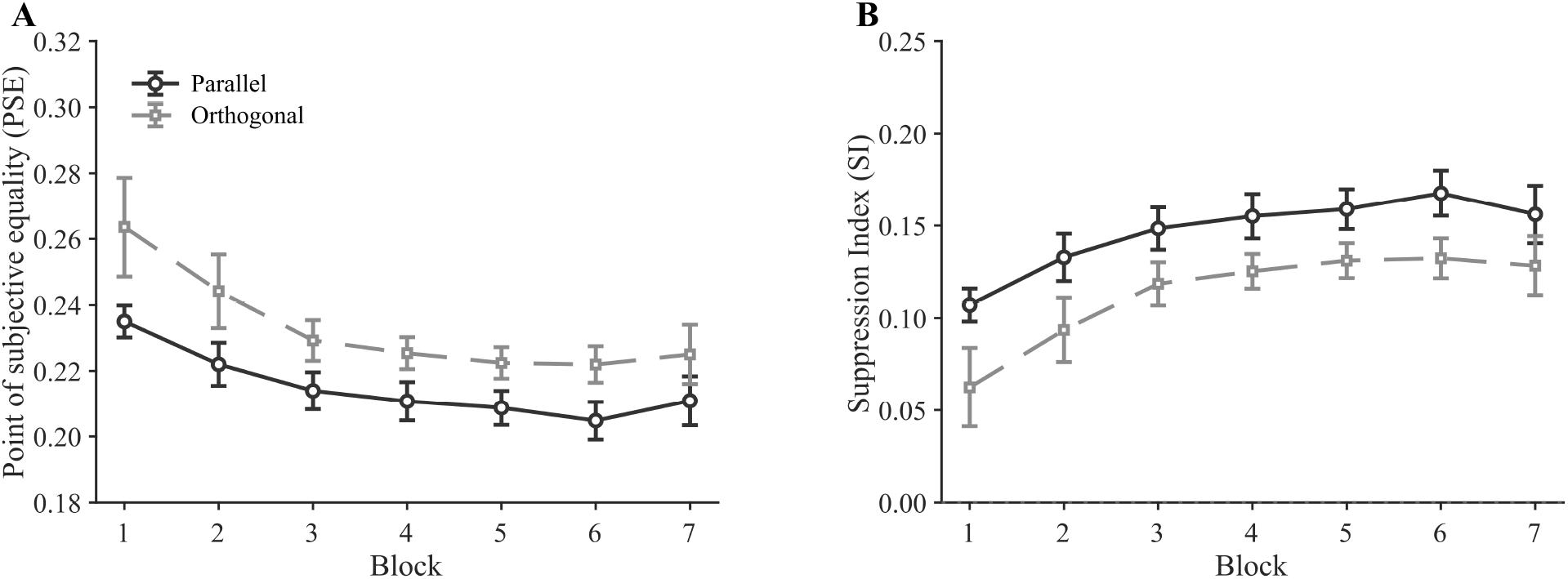
Results in Experiment 1. *Note.* (A) Mean PSE for the central grating in the parallel and orthogonal conditions. (B) Mean SI. Higher values indicate stronger contrast surround suppression. Error bars represent ±1 *SEM*.

In summary, Experiment 1 showed that, during repeated measurement blocks using static center-surround stimuli, contrast surround suppression increased after the first measurement block and remained elevated across most subsequent blocks. We found no evidence that this change could be modulated by the relative orientation of the center and surround. Thus, contrast surround suppression varied on the timescale of minutes as a function of recent center-surround stimulus experience.

## Experiment 2

Experiment 1 showed that contrast surround suppression changed with short-term stimulus history. Experiment 2 tested whether a comparable effect generalized to motion surround suppression. We predicted that motion surround suppression would strengthen across repeated measurement blocks that contained repeated presentation of large drifting gratings.

### Method

#### Participants

As in Experiment 1, sample-size planning was based on repeated-measures ANOVA with seven time points (see Data Analysis), assuming a large effect size (*η*^2^ = .14), *α* = .05, and power (1 − *β*) = .80. This yielded a minimum sample size of 8 participants. We recruited 14 students from Zhejiang University, including one author. All participants except the author were naive to the purpose of the experiment. All 14 were included in the final analysis (9 women, 5 men; age range = 18–22 years, *M* = 19.4, *SD* = 1.0). Participants provided written informed consent and received either monetary compensation or course credit. All procedures were approved by the Research Ethics Committee of Zhejiang University.

#### Apparatus and Stimuli

Visual stimuli were generated in MATLAB R2014a using Psychtoolbox (Brainard, 1997; Pelli, 1997) and presented on a 32-in. Display++ monitor (Cambridge Research Systems Ltd., Rochester, UK; resolution = 1,920 × 1,080 pixels; refresh rate = 100 Hz). The monitor was automatically gamma-linearized, and mean background luminance was 40.26 cd/m². Participants viewed the stimuli binocularly in a dimly lit room at a fixed viewing distance of 75 cm, maintained with a chinrest.

Experiment 2 used high-contrast small and large drifting gratings (Figure 3A), subtending 1° and 5° of visual angle in diameter, respectively. The outermost 0.3° of each grating was tapered with a cosine envelope. Apart from size, the two gratings were identical: Michelson contrast = 0.90, drift speed = 3.5°/s, spatial frequency = 1.5 cpd, and horizontal orientation. On each trial, the grating drifted vertically upward or downward randomly with equal probability. Stimulus contrast was modulated by a Gaussian temporal envelope truncated at ±4σ (total duration = 8σ). For subsequent analyses, stimulus duration was defined as 2σ of this temporal envelope.

**Figure 3.**
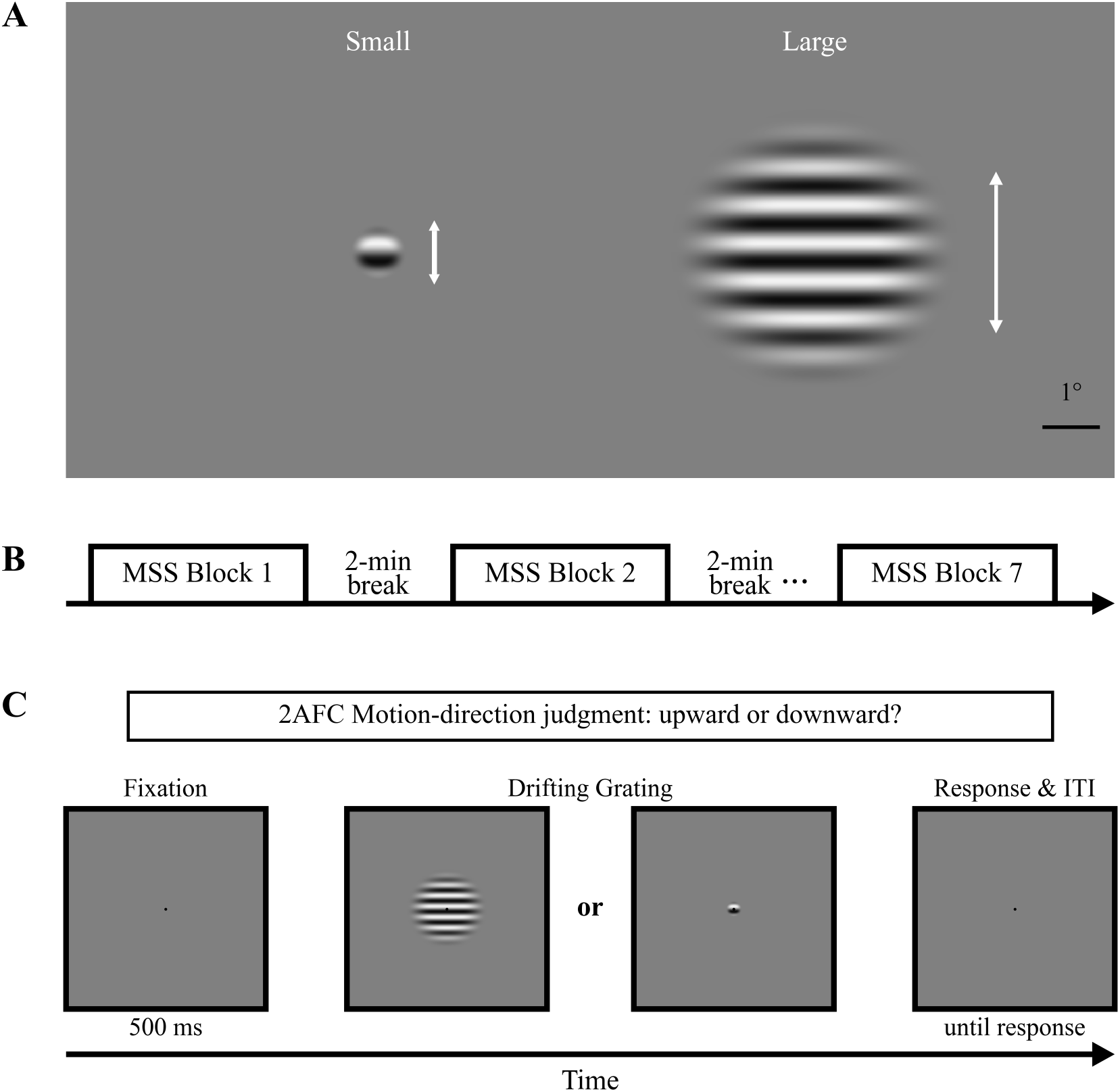
Stimuli and Procedure in Experiment 2. *Note.* (A) Small and large drifting gratings used in Experiment 2. The gratings moved upward or downward with equal probability. (B) Session structure. The session contained seven 280-trial measurement blocks. Participants took a 30-s break halfway through each block and a 1-min break between blocks. (C) Trial sequence. Following fixation, a small or large drifting grating was presented at the center of the screen. Participants reported whether the grating moved upward or downward. Stimuli are shown for illustration and are not drawn to scale. 2AFC = two-alternative forced choice.

#### Procedure

As shown in Figure 3B, Experiment 2 comprised seven consecutive measurement blocks of 280 trials each. Participants took a 30-s break after the 140th trial within each block and a 1-min break between blocks. During all breaks, they were instructed to close their eyes to minimize additional visual input. The experiment comprised 1,960 trials and lasted approximately 1 hr. Before formal testing, participants completed practice trials with the small grating to familiarize themselves with the task. Practice data were not analyzed.

Motion surround suppression was measured with a duration-threshold paradigm using a two-alternative forced-choice task (Figure 3C) (Tadin et al., 2003). Each trial began with a 500-ms fixation display, followed by a small or large drifting grating at fixation. The grating moved upward or downward randomly with equal probability, with stimulus size and motion direction randomized across trials. After stimulus offset, participants reported the motion direction by keypress. Incorrect responses were followed by auditory feedback, while correct responses received no feedback. Within each block, four randomly interleaved 1-up-3-down staircases controlled stimulus duration for the small and large gratings. Each size was assigned two 70-trial staircases, yielding 140 trials per size and 280 trials per block. For each size, the two staircases began at 5 and 120 ms. Stimulus duration was selected from 13 logarithmically spaced levels between 5 and 120 ms. Following an incorrect response, duration increased by one level the next time that staircase was sampled, while after three consecutive correct responses within a staircase, duration decreased by one level. Motion surround suppression was then estimated by fitting psychometric functions to all trials within each block.

#### Data Analysis

For each repeated measurement block and each participant, the 140 trials from the two staircases corresponding to each grating size were pooled separately for psychometric fitting. At each duration level *D*, we calculated motion-direction discrimination accuracy, *P*(*D*). Using MATLAB’s fitnlm function, accuracy was then fitted with a cumulative Gaussian psychometric function incorporating a 50% guessing rate:

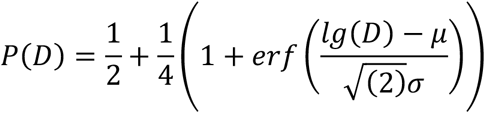

Here, *μ* and *σ* denote the location and scale parameters, respectively, of the cumulative Gaussian function on the log-duration axis. The duration at which the fitted function reached *P*(*D*) = .7937 was defined as the motion-direction discrimination threshold. Each participant therefore yielded one duration threshold for the small grating and one for the large grating in each block.

We then defined the Suppression Index (SI) as lg(TL/TS), where TL and TS denote the motion-direction discrimination duration thresholds for the large and small gratings, respectively. Thus, SI > 0 indicates a higher threshold for the large than for the small grating, suggesting poorer motion-direction discrimination with increasing stimulus size. Therefore, larger SI values indicate stronger motion surround suppression.

To test whether motion surround suppression changed with repeated measurement, SI was submitted to a one-way repeated-measures ANOVA with Block (1–7) as a within-participant factor. Sphericity was assessed with Mauchly’s test, and Greenhouse-Geisser corrections were applied when the assumption was violated. If the main effect of Block was significant, follow-up pairwise comparisons were Bonferroni-corrected. The significance criterion was α = .05. All statistical analyses were conducted in JASP 0.95.4 (JASP Team, 2025)

#### Results

Figure 4B shows SI across repeated measurement blocks in Experiment 2. A one-way repeated-measures ANOVA with Block (1–7) as a within-participant factor revealed no significant main effect of Block, *F*(2.53, 32.88) = .30, *p* = .79, *η*^2^ = .023. Thus, we did not detect a systematic change in motion surround suppression across repeated measurement blocks.

**Figure 4.**
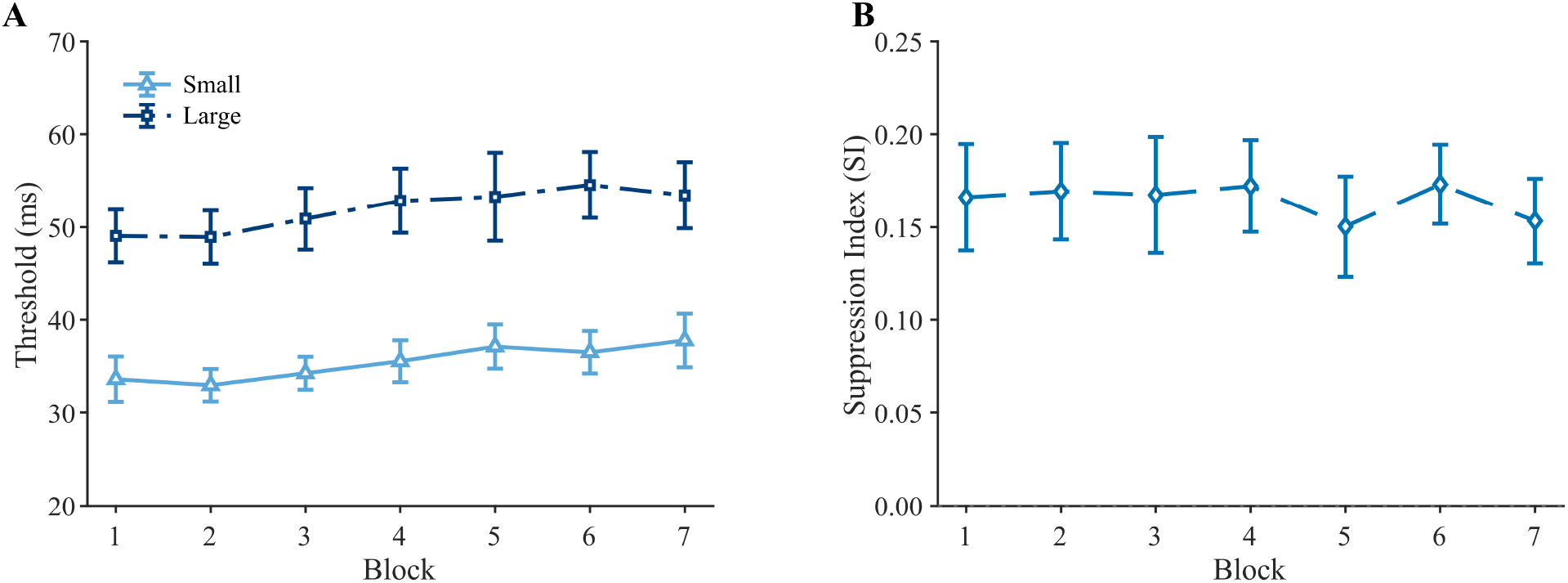
Results in Experiment 2. *Note.* (A) Mean motion-direction discrimination thresholds for the small and large gratings across blocks. (B) Mean SI. Higher values indicate stronger motion surround suppression. Error bars represent ±1 *SEM*.

## Experiment 3

Experiment 1 showed that contrast surround suppression changed with short-term stimulus history, whereas Experiment 2 did not reveal a comparable change in motion surround suppression. This difference raises the possibility that the spatiotemporal properties of the visual stimulus constrain the time course of suppression change. Experiment 3 therefore tested whether short-term changes in contrast surround suppression depend on stimulus spatiotemporal properties. We predicted that the static, high-spatial-frequency condition would replicate the increase in suppression observed in Experiment 1, whereas the accumulation of suppression across blocks would be weaker or less apparent for the dynamic, low-spatial-frequency flickering stimuli.

### Method

#### Participants

As in Experiment 1, sample-size planning was based on repeated-measures ANOVA with seven time points (see Data Analysis), assuming a large effect size (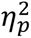 = .14), *α* = .05, and power (1 − *β*) = .80. This yielded a minimum sample size of 8 participants. We recruited 16 students from Zhejiang University, including one author. All participants except the author were naive to the purpose of the experiment. All 16 were included in the final analysis (7 women, 9 men; age range = 18–22 years, *M* = 19.6, *SD* = 1.3). Participants provided written informed consent and received either monetary compensation or course credit. All procedures were approved by the Research Ethics Committee of Zhejiang University.

#### Apparatus and Stimuli

Visual stimuli were generated in MATLAB R2014a using Psychtoolbox (Brainard, 1997; Pelli, 1997) and presented on a 32-in. Display++ monitor (Cambridge Research Systems Ltd., Rochester, UK; resolution = 1,920 × 1,080 pixels; refresh rate = 100 Hz). The monitor was automatically gamma-linearized. Mean background luminance was 40.26 cd/m². Participants viewed the stimuli binocularly in a dimly lit room at a fixed viewing distance of 75 cm, maintained with a chinrest.

Experiment 3 included two stimulus conditions, static and dynamic, illustrated in Figure 5. In both conditions, the standard stimulus was a center-surround grating and the matching stimulus was the central circular grating presented alone. Michelson contrast of the center and surround in the standard stimulus was fixed at 0.30 and 0.80, respectively, whereas matching-stimulus contrast was controlled by the staircase. The static condition used exactly the same stimulus parameters as the parallel condition in Experiment 1. In the dynamic condition, spatial frequency was reduced to 1 cpd for both standard and matching gratings, and the gratings counterphase flickered at 8 Hz. All other parameters were identical to those in the static condition. In both conditions, grating orientation was randomly set on each trial to 45° clockwise or counterclockwise from vertical. The surround was always parallel to the center in the standard stimulus. Stimulus onset and offset were each tapered with a 100-ms cosine ramp.

**Figure 5.**
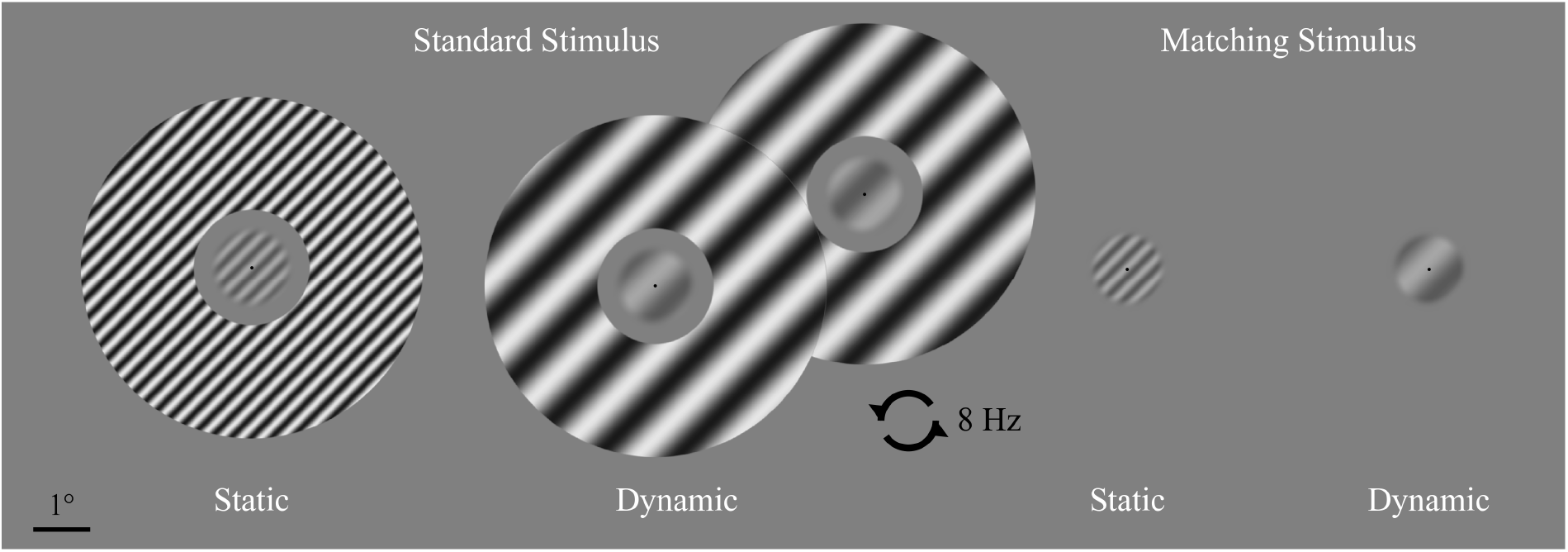
Stimuli and Procedure in Experiment 3. *Note.* Standard and matching stimuli in the static and dynamic conditions. The static condition used the same center-surround configuration as the parallel condition in Experiment 1. The dynamic condition used lower-spatial-frequency gratings that counterphase flickered at 8 Hz.

#### Procedure

The static and dynamic conditions were completed on separate testing days at least 24 hr apart, with condition order counterbalanced across participants. The task and session structure were otherwise identical to Experiment 1 (Figures 1B–1C), with seven measurement blocks per condition. Each block comprised 160 trials, with a 30-s break after Trial 80 and a 2-min break between adjacent blocks. During breaks, participants were instructed to close their eyes to minimize additional visual input. Each condition comprised 1,120 trials and lasted approximately 1.5 hr. On the day before formal testing, participants practiced both stimulus conditions; practice data were not analyzed.

#### Data Analysis

For each repeated measurement block in each stimulus condition for every participant, psychometric functions were fitted and SI was calculated using the same procedures as in Experiment 1 (see Experiment 1, Data Analysis). To compare the short-term change in contrast surround suppression between the static and dynamic conditions, SI was submitted to a two-way repeated-measures ANOVA with Stimulus Condition (static, dynamic) and Block (1–7) as within-participant factors. Sphericity was assessed with Mauchly’s test, and Greenhouse-Geisser corrections were applied when the assumption was violated. Bonferroni correction was used for all possible post hoc comparisons. All statistical analyses were conducted in JASP 0.95.4 (JASP Team, 2025).

#### Results

Figure 6 shows PSE and SI across blocks in the static and dynamic conditions. A two-way repeated-measures ANOVA on SI revealed a significant main effect of Stimulus Condition, *F*(1, 15) = 12.99, *p* = .003, 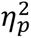 = .46, and a significant main effect of Block, *F*(1.47, 22.05) = 5.03, *p* = .024, 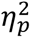 =.25. Critically, the Stimulus Condition × Block interaction was also significant, *F*(2.18, 32.65) = 7.99, *p =* .001, 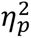 =.35. We therefore examined the simple effect of Block within each condition. In the static condition, SI varied significantly across blocks, *F*(1.47, 22.00) = 10.34, *p =* .002, *η*^2^ =.41. Bonferroni-corrected comparisons showed that SI in Block 1 was lower than in Block 2 (adjusted *p* < .01) and lower than in Blocks 3, 6, and 7 (all adjusted *p*s < .05). Thus, in the static condition, SI increased after the first block and remained elevated in several subsequent blocks. By contrast, SI did not vary significantly across blocks in the dynamic condition, *F*(1.85, 27.75) = .48, *p =* .61, *η*^2^ =.03. Experiment 3 therefore did not detect a systematic increase in contrast surround suppression across blocks under dynamic stimulation.

**Figure 6.**
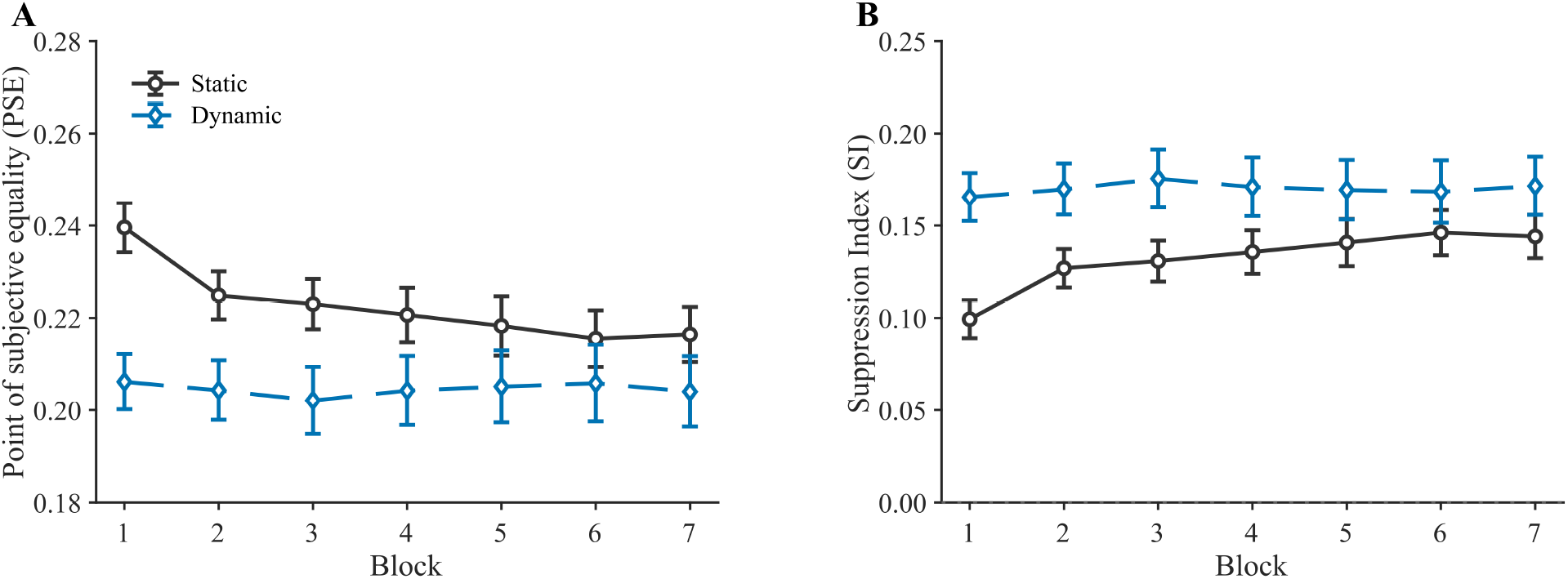
Results in Experiment 3. *Note.* (A) Mean PSE for the central grating in the static and dynamic conditions. (B) Mean SI. Higher values indicate stronger contrast surround suppression. Error bars represent ±1 *SEM*.

In summary, Experiment 3 revealed an increase in contrast surround suppression only in the static condition but not in the dynamic condition. One possibility is that suppression may rise too rapidly to be detected by the current procedural setup. Experiment 4 was therefore designed to test this possibility by lengthening each block and removing within-block breaks, providing a higher temporal resolution.

## Experiment 4

In Experiment 3, contrast surround suppression increased after the first measurement block in the static condition, whereas no systematic across-block change was observed in the dynamic condition. One possibility is that dynamic stimulation may induce a faster suppression change. To sample the time course with greater sensitivity, Experiment 4 lengthened the measurement blocks and eliminated within-block breaks, creating a more sustained and stable center-surround stimulation environment. We predicted an increase in both conditions, with the dynamic condition potentially reaching an asymptote more rapidly.

### Method

#### Participants

As in the preceding experiments, sample-size planning was based on a paired-samples t test (see Data Analysis), assuming a conventionally large effect size (Cohen’s *d* = .80), *α* = .05, and power (1 − *β*) = .80. This yielded a minimum sample size of 15 participants. Experiment 4 recruited 21 students from Zhejiang University, all of whom were included in the final analysis (12 women, 9 men; age range = 18–25 years, *M* = 19.1, *SD* = 1.6). All participants were right-handed, had normal or corrected-to-normal vision, and reported no history of psychiatric illness. Participants provided written informed consent and received either monetary compensation or course credit. All procedures were approved by the Research Ethics Committee of Zhejiang University.

#### Apparatus and Stimuli

The hardware and software setup in Experiment 4 was identical to that used in Experiment 3. The static and dynamic stimulus parameters were also identical to those in Experiment 3 (see Experiment 3, Apparatus and Stimuli, and Figure 5).

#### Procedure

The static and dynamic conditions were completed on separate testing days at least 16 hr apart, with condition order counterbalanced across participants. Each condition comprised three consecutive measurement blocks of 240 trials, with no breaks within a block and a 150-s break between blocks. During between-block breaks, participants were instructed to close their eyes to minimize additional visual input. Each trial used the same 2IFC contrast-matching task as in Experiments 1 and 3. Each condition comprised 720 trials and lasted approximately 50 min. On the day before formal testing, participants practiced both stimulus conditions. Practice data were not analyzed.

#### Data Analysis

For each participant and stimulus condition, SI was calculated in the same manner as in Experiments 1 and 3. Larger SI values indicate stronger contrast surround suppression.

The primary aim of Experiment 4 was to determine whether suppression increased in both stimulus conditions. We reasoned that the increase might occur rapidly early in the session and then approach an asymptote, a pattern also suggested by the static conditions in Experiments 1 and 3. Experiment 4 therefore first tested for the suppression increase within Block 1. Specifically, SI values calculated from the first and final 80 trials were compared using one-tailed paired-samples *t* tests, separately for the static and dynamic conditions. Experiment 4 also used an 80-trial sliding window with a 40-trial step to obtain five overlapping windows within Block 1. For each participant and condition, a linear function was fitted to the resulting SI series, and the mean slope was tested against zero using a one-tailed one-sample *t* test to assess whether SI showed an overall positive trend across Block 1.

Also, Experiment 4 examined suppression change across the full experiment. To characterize the trajectory over the session, we applied the same 80-trial window with a 40-trial step to all 720 temporally ordered trials, yielding 17 overlapping windows. For each participant and condition, the SI series was min-max normalized to the range 0-1 and fitted with the following saturating exponential function:

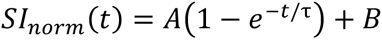

In this model, *A* represents the change in suppression from baseline to asymptote, *B* represents the baseline, and *τ* is the time constant. Smaller *τ* values indicate a faster model-estimated temporal change. To assess suppression change across the full experiment at the group level, we resampled participants with replacement 5,000 times. For each bootstrap sample, we calculated the group-mean SI trajectory and fitted the exponential model. This procedure yielded a 95% one-sided lower bootstrap confidence bound for the change-amplitude parameter A, separately for the static and dynamic conditions. An increase in suppression was inferred when the lower confidence bound exceeded zero.

The second aim of Experiment 4 was to test whether suppression increased more rapidly in the dynamic than in the static condition. To this end, Experiment 4 examined the time constant *τ* from the exponential model in both the participant-level and group-mean-level analyses. Because estimates of *τ* are interpretable only when the underlying series shows a reliable change in suppression, the participant-level analysis first used bootstrap resampling to identify participants who showed reliable suppression increases in both conditions. Specifically, for each participant and condition, the first 120 and last 120 trials were each resampled with replacement 5,000 times, and the late-minus-early SI difference was computed for every resample. A participant was included when the 95% one-sided lower bootstrap confidence bound for this difference exceeded zero in both conditions. The exponential model was then fitted at the individual-participant level only for participants who met this criterion in both conditions. Their time constants were compared using a one-tailed paired-samples *t* test to test whether *τ* was smaller in the dynamic than in the static condition.

At the group-mean level, participants were first resampled with replacement 5,000 times. For each bootstrap sample, the group-mean trajectories were calculated and fitted with the exponential model, yielding a bootstrap distribution of the between-condition difference in time constants, Δ*τ* (dynamic − static). A 95% one-sided upper bootstrap confidence bound was obtained for Δ*τ*, and a faster change in the dynamic condition was inferred when this upper bound was below zero. Experiment 4 also tested the time-constant difference with a nonparametric permutation procedure. Specifically, condition labels were randomly exchanged within participants for 5,000 permutations. On each permutation, group-mean trajectories were calculated, the exponential model was fitted, and Δ*τ* was obtained as the test statistic. These Δ*τ* values therefore formed a permutation null distribution. The one-tailed permutation *p* value was the proportion of the Δ*τ* values that were smaller than the observed one, with *α* = .05. All analyses were performed in MATLAB R2022b.

#### Results

To determine whether suppression increased in both stimulus conditions, we first focused on the early part of the session because Experiments 1 and 3 suggested an initial increase in suppression followed by an asymptote. Figure 7A shows SI across Block 1 in Experiment 4. In the static condition, SI was significantly higher in the final 80 trials than in the first 80 trials (*M* = .020, *SD* = .027), *t*(20) = 3.45, *p* = .001, Cohen’s *d* = .75. The same pattern was observed in the dynamic condition (*M* = .025, *SD* = .041), *t*(20) = 2.80, *p* = .006, Cohen’s *d* = .61 (Figure 7B). To assess whether suppression increased progressively within Block 1, we fitted a linear function to SI across the five sliding windows. Mean slopes were significantly greater than zero in both the static condition (*M* = .0058, *SD* = .0067), *t*(20) = 3.95, *p* < .001, Cohen’s *d* = .86, and the dynamic condition (*M* = .0062, *SD* = .012), *t*(20) = 2.46, *p* = .012, Cohen’s *d* = .54 (Figure 7C). These results suggested that contrast surround suppression increased within the first block in both conditions.

**Figure 7.**
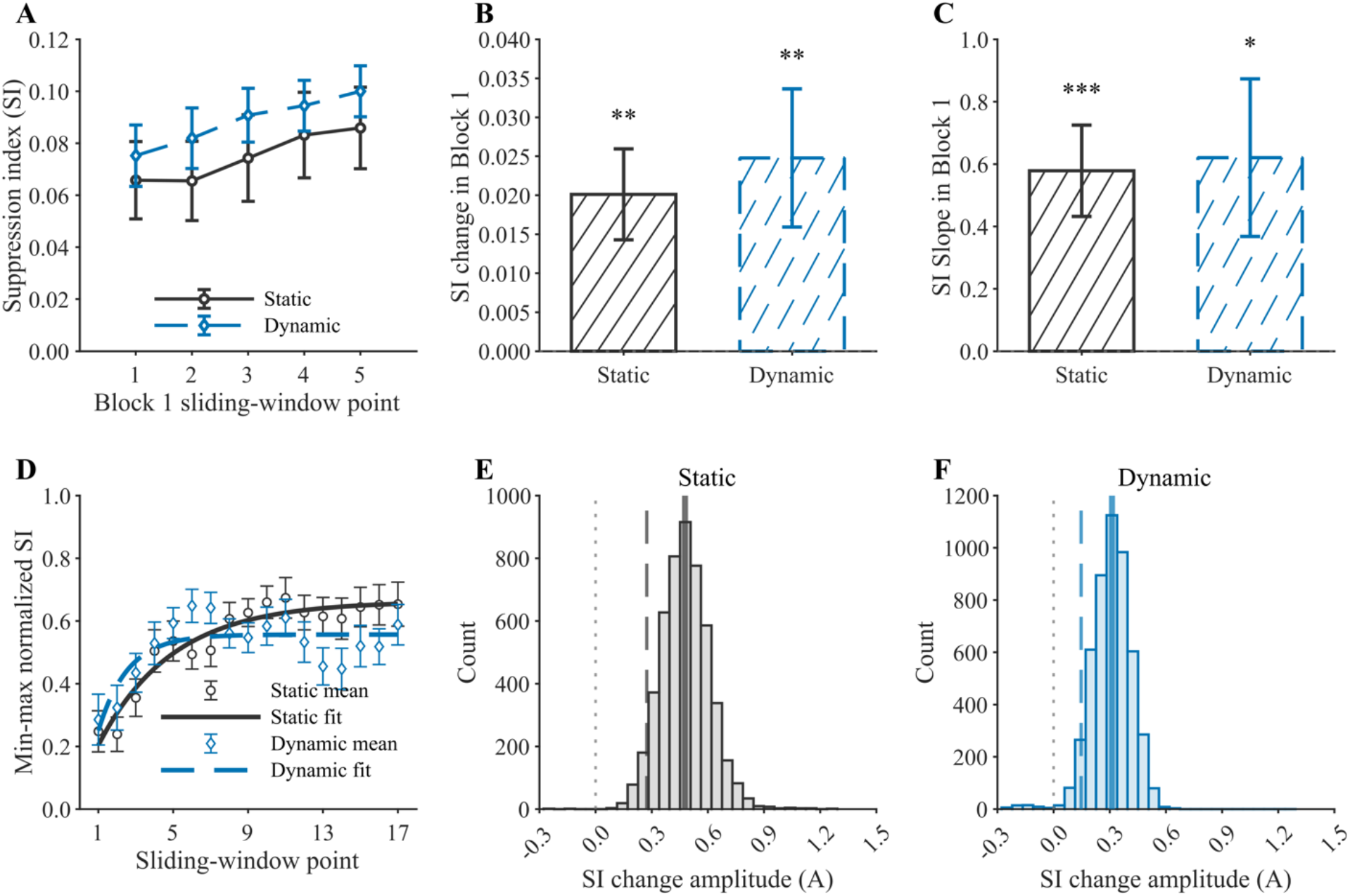
Results in Experiment 4. *Note.* (A) Mean SI across five overlapping windows in Block 1 (80-trial window, 40-trial step). Dark open circles indicate the static condition, and light-blue diamonds indicate the dynamic condition. (B) Change in SI from the first to the last 80-trial segment of Block 1. (C) Participant-level linear slopes fitted across the five Block 1 windows. (D) Group-mean min-max-normalized SI trajectories and saturating exponential fits. (E–F) Bootstrap distributions of the change-amplitude parameter A for the static and dynamic conditions. Solid lines indicate bootstrap medians, dark long-dashed lines indicate the 95% one-sided lower bootstrap confidence bound, and light short-dashed lines mark zero. Error bars represent ±1 SEM. \**p* < .05. \*\**p* < .01. \*\*\**p* < .001.

We next examined suppression change across the full session by fitting the saturating exponential model to the group-mean normalized SI trajectories (Figure 7D). For the static condition, the estimated change-amplitude parameter was *A* = .46, with a 95% one-sided lower bootstrap confidence bound of .27. For the dynamic condition, the corresponding estimate was *A* = .31, with a lower bound of .15. Both lower bounds were above zero (Figures 7E and 7F), providing evidence that contrast surround suppression increased over the course of the whole experiment in both stimulus conditions. Together with the Block 1 analyses, these results indicate that contrast surround suppression strengthened in both the static and dynamic conditions.

We next compared the rate of suppression change between the static and dynamic conditions by analyzing the time constant *τ* of the saturating exponential model at the individual and group levels. Because *τ* can be estimated reliably only when the underlying series shows a clear suppression change, we first used bootstrap resampling to identify participants with reliable increases across the full experiment (Figure 8A). Of the 21 participants, 15 showed a reliable increase in the static condition, 9 in the dynamic condition, and 7 in both conditions. We then fitted the saturating exponential model separately to each of these seven participants. Their time constants were smaller in the dynamic condition (*M* = 3.15, *SD* = 1.83) than in the static condition (*M* = 10.79, *SD* = 8.81), *t*(6) = −2.37, *p* = .028, Cohen’s *d* = −.90 (Figure 8B). Thus, among participants who showed a reliable suppression increase in both conditions, the increase was faster under the dynamic condition.

**Figure 8.**
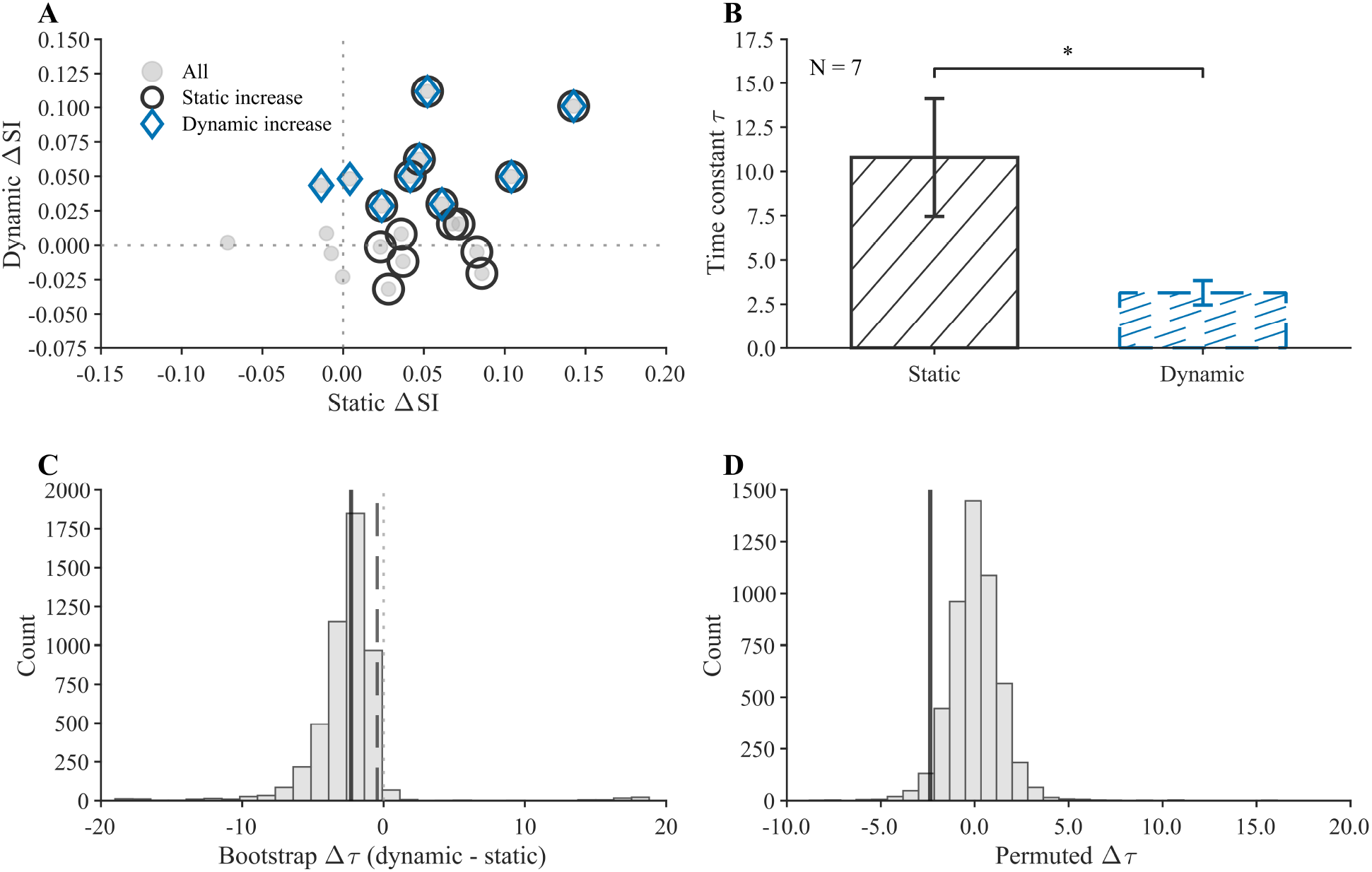
Comparison of Suppression-Change Time Courses in Experiment 4. *Note.* (A) Participant-level bootstrap classification of early-to-late SI change in the static and dynamic conditions. The x-and y-axes show suppression change in the static and dynamic conditions, respectively. Gray circles represent individual participants; black circles identify participants with a reliable increase in the static condition, and blue diamonds identify participants with a reliable increase in the dynamic condition. (B) Time constants for participants who showed reliable increases in both conditions (*n* = 7). (C) Group-level bootstrap distribution of Δ*τ*. The solid line indicates the bootstrap median, dark long-dashed lines indicate the 95% one-sided upper confidence bound, and the light short-dashed line marks zero. (D) Within-participant permutation null distribution of Δ*τ*. The solid line marks the observed value. Error bars represent ±1 *SEM*. \**p* < .05.

At the group-mean level, the observed time-constant difference was Δ*τ* (dynamic − static) = −2.37. Bootstrap resampling yielded a 95% one-sided upper confidence bound of - 0.44 (Figure 8C), consistent with a smaller time constant in the dynamic condition. Furthermore, the nonparametric permutation test showed that the observed difference was significantly smaller than expected under the permutation null distribution, *p* = .030 (Figure 8D). Together with the participant-level analysis, these results suggest that contrast surround suppression increased more rapidly under dynamic than under static stimulation.

In summary, building on Experiment 3, Experiment 4 showed that contrast surround suppression strengthened in both static and dynamic conditions, but the increase may have occurred more rapidly under dynamic stimulation. This pattern points to stimulus-dependent differences in the short-term plasticity of surround suppression.

## Discussion

The present study systematically examined whether perceptual surround suppression changes with recent visual experience during repeated measurement. Across four psychophysical experiments, three main findings emerged. First, with static center-surround stimuli, contrast surround suppression increased early during repeated measurement and then approached a plateau. Second, with dynamic stimuli characterized by lower spatial and higher temporal frequencies, an increase in contrast surround suppression was also observed, but the change may have occurred more rapidly than under static stimulation. Third, with drifting gratings, we found no systematic change in motion surround suppression across repeated measurement. Taken together, these findings indicate that contrast surround suppression is not completely stable within a testing session. It can change over minutes as a function of recent stimulus experience, and the time course of that change may depend on stimulus properties. At the same time, we found no evidence that this short-term change generalizes to motion surround suppression.

These findings have implications for how contrast surround suppression is conceptualized and measured. Psychophysical studies commonly pool large numbers of trials within a condition and summarize suppression with a single PSE, threshold, or suppression index (Cai et al., 2008; Xing & Heeger, 2001; Zenger-Landolt & Heeger, 2003). Across three experiments with almost entirely independent samples, however, we repeatedly observed a clear early increase in contrast surround suppression under static stimulation. The finer temporal sampling in Experiment 4 further showed that this increase continued across the first 240 trials before remaining elevated and gradually approaching an asymptote. At least under the present stimulus and task conditions, surround-suppression strength is therefore better viewed as a dynamic perceptual state shaped jointly by current stimulus properties and recent stimulus history, rather than as a completely fixed measurement parameter. This also means that when suppression is estimated from long testing sequences, the visual experience accumulated during the measurement itself may contribute to the resulting estimate.

The experience-dependent change observed here is consistent with the dynamic normalization account formalized in the Hebbian normalization model (Westrick et al., 2016). In classical divisive-normalization models, responses of a target neural population are regulated by the weighted activity of a normalization pool (Carandini & Heeger, 2012). The Hebbian normalization model extends this framework by allowing the normalization weights to change dynamically with recent coactivation following Hebbian rules. When two interacting populations frequently respond together, the normalization weight between them increases, strengthening their mutual suppressive influence and thereby reducing subsequent coactivation and representational redundancy. This account was first supported by the neurophysiological findings of Aschner et al. (2018), who found that adaptation to two spatially overlapping gratings presented synchronously increased subsequent cross-orientation suppression, whereas alternating presentation of the same gratings reduced it. Yiltiz et al. (2020) reported a similar pattern in human psychophysics, and the Hebbian normalization model captured both the electrophysiological and psychophysical changes. More recent EEG and MEG evidence further suggests that normalization reweighting can emerge within seconds of sustained stimulation (Baker et al., 2023). Because divisive normalization is widely considered a computational mechanism underlying contrast surround suppression (Cavanaugh et al., 2002; Xing & Heeger, 2001), one possibility is that repeated center-surround presentation in our experiments repeatedly coactivated center- and surround-related neural populations, strengthening their normalizing interaction and thereby increasing perceptual surround suppression. Our findings are therefore compatible with the view that visual normalization is not entirely fixed. They further show that measurable changes in surround suppression can arise from the repeated center-surround relationships encountered during an ordinary psychophysical measurement sequence, without a separately imposed prolonged adaptation phase.

Experiment 1 further suggests that the stimulus factors determining the overall magnitude of contrast surround suppression may not identically govern its short-term change. Consistent with previous work (Xing & Heeger, 2001), overall suppression was substantially stronger when center and surround were parallel than when they were orthogonal. However, we found no evidence that the short-term increase in suppression depended on this relative orientation. This pattern resembles that reported by Yiltiz et al. (2020), who observed increased contrast surround suppression after center-surround adaptation for both 30° and 90° orientation differences, with no significant difference in the magnitude of the increase. Neural accounts of contrast surround suppression have implicated surround suppression in the primary visual cortex (V1), including early and late components with different tuning properties (Schallmo & Murray, 2016; Schallmo et al., 2019; Webb et al., 2005). The early component is monocular and relatively insensitive to orientation, whereas the later component is binocular and strongly orientation selective, with maximal suppression when center and surround are parallel. The orientation-invariant plasticity observed here may therefore partly reflect an enhancement of an earlier, broadly tuned component of contrast surround suppression.

Experiments 3 and 4 further showed that the observability and time course of short-term changes in contrast surround suppression depend on the spatiotemporal properties of the stimulus and on how suppression is sampled. Experiment 3 replicated the clear across-block suppression increase observed in Experiment 1 under static stimulation, but did not detect the same pattern with low-spatial-frequency counterphase-flickering stimuli. When Experiment 4 extended the uninterrupted measurement periods and increased temporal sampling resolution early in the session, however, we observed that suppression increased within the first block in both the static and dynamic conditions, and the saturating exponential model suggested a smaller time constant under dynamic stimulation. Considered together, the two experiments suggest that detecting suppression change under dynamic stimulation may require a more sustained and stable history of center-surround input, and that the change itself may unfold more rapidly. One speculative possibility concerns differences between magnocellular and parvocellular processing. The static stimuli used in Experiments 3 and 4 may preferentially engage processing sensitive to higher spatial and lower temporal frequencies, often associated with the parvocellular (P) pathway, whereas the dynamic stimuli may preferentially engage processing sensitive to lower spatial and higher temporal frequencies, often associated with the magnocellular (M) pathway (Solomon, 2021). Differences in short-term plasticity between the two stimulus conditions could therefore partly reflect differences in the plasticity of *M*- and P-related processing. Interestingly, short-term monocular deprivation studies provide a suggestive parallel (Binda et al., 2018). After brief monocular deprivation, perceptual dominance of the deprived eye transiently increases (Lunghi et al., 2011), an effect commonly interpreted as compensatory homeostatic plasticity (Turrigiano, 2011). Binda et al. (2018) further reported that this effect was more closely associated with higher spatial frequencies and the ventral visual pathway, leading them to propose stronger short-term plasticity in P-pathway-related processing.

In contrast to contrast surround suppression, Experiment 2 did not reveal a systematic change in motion surround suppression across repeated measurement. This raises the possibility that, on the timescale of several minutes, dynamic changes in motion surround suppression may require finer temporal sampling or a more sustained stimulation context to be detected. Although contrast surround suppression and motion surround suppression both reflect contextual suppression of local visual processing and can both be captured by normalization-based accounts (Cavanaugh et al., 2002; Schallmo et al., 2018; Xing & Heeger, 2001), they are thought to rely on partly different neural mechanisms. Contrast surround suppression has been linked primarily to neural surround suppression in human V1, whereas motion surround suppression has been linked more closely to suppression in MT/hMT+ (Schallmo et al., 2018; Zenger-Landolt & Heeger, 2003). The different patterns observed here across stimulus history may therefore reflect differences in short-term plasticity across visual cortical regions. Consistent with this possibility, Binda et al. (2018) found no comparable short-term monocular-deprivation effect in hMT+, again suggesting regional differences in homeostatic plasticity. Motion surround suppression may therefore differ from contrast surround suppression in its short-term plasticity, but this possibility requires more systematic investigation.

One may argue that several alternative explanations for the block-related increase in contrast surround suppression should be considered, including fatigue, task familiarization, decision drift, and visual adaptation. Several aspects of the design and results make it unlikely that any one of these factors was the dominant source of the effect. First, all participants completed substantial practice on the day before formal testing, reducing the likelihood that the observed change primarily reflected early task familiarization. Second, a uniform fatigue effect or decision drift would have difficulty accounting for the different temporal profiles observed across stimulus conditions in Experiments 3 and 4. Finally, we cannot exclude a contribution of visual adaptation during repeated measurement, but ordinary adaptation alone does not readily predict the observed direction of change. Both center-alone and center-surround stimuli were presented repeatedly, and if adaptation weakened the perceived central stimulus in both contexts and thereby reduced their perceptual difference, the expected consequence would be weaker, not stronger, contrast surround suppression. We therefore consider the observed pattern more likely to reflect an increase in contrast surround suppression itself than a nonspecific consequence of fatigue, practice, decision drift, or conventional visual adaptation.

The present study also has several limitations that motivate further work. First, we did not directly manipulate the temporal contingency between center and surround and therefore cannot determine whether the increase in suppression shows the correlation dependence predicted by the Hebbian normalization model. Future studies could hold overall center and surround stimulation constant while systematically comparing synchronous, temporally decorrelated, and asynchronous center-surround sequences, asking whether suppression changes in the direction predicted by their joint statistics. Computational modeling could then provide a more direct test of whether the observed change conforms to the Hebbian normalization framework and help constrain the underlying computation. Second, the static and dynamic conditions in Experiments 3 and 4 differed simultaneously in spatial and temporal frequency. The specific contribution of each dimension to short-term plasticity of contrast surround suppression therefore remains unresolved. Orthogonal manipulation of spatial and temporal frequency will be needed to determine how each shapes the time course of suppression change. More broadly, future work could systematically compare the plasticity of different visual pathways and cortical regions. Finally, although we did not observe a comparable change in motion surround suppression, Experiment 2 did not sample its time course at the finer temporal resolution used in Experiment 4. More temporally resolved and systematic measurements will therefore be needed to characterize any possible short-term plasticity of motion surround suppression.

## Conclusion

Taken together, the present study shows that perceptual contrast surround suppression can change within a single testing session, increasing early and then approaching a plateau, with the observability of this change depending on stimulus and sampling conditions. The findings are compatible with rapid, experience-dependent reweighting of contextual normalization. The absence of a comparable change in motion surround suppression further indicates that short-term plasticity should not be assumed to generalize across behavioral phenomena that share the label of surround suppression.

